# ChIP-exo interrogation of Crp, DNA, and RNAP holoenzyme interactions

**DOI:** 10.1101/069021

**Authors:** Haythem Latif, Stephen Federowicz, Ali Ebrahim, Janna Tarasova, Richard Szubin, Jose Utrilla, Karsten Zengler, Bernhard O. Palsson

## Abstract

Numerous *in vitro* studies have yielded a refined picture of the structural and molecular associations between Cyclic-AMP receptor protein (Crp), the DNA motif, and RNA polymerase (RNAP) holoenzyme. In this study, high-resolution ChIP-exonuclease (ChIP-exo) was applied to study Crp binding *in vivo* and at genome-scale. Surprisingly, Crp was found to provide little to no protection of the DNA motif under activating conditions. Instead, Crp demonstrated binding patterns that closely resembled those generated by σ^70^. The binding patterns of both Crp and σ^70^ are indicative of RNAP holoenzyme DNA footprinting profiles associated with stages during transcription initiation that occur post-recruitment. This is marked by a pronounced advancement of the template strand footprint profile to the +20 position relative to the transcription start site and a multimodal distribution on the nontemplate strand. This trend was also observed in the familial transcription factor, Fnr, but full protection of the motif was seen in the repressor ArcA. Given the time-scale of ChIP studies and that the rate-limiting step in transcription initiation is typically post recruitment, we propose a hypothesis where Crp is absent from the DNA motif but remains associated with RNAP holoenzyme post-recruitment during transcription initiation. The release of Crp from the DNA motif may be a result of energetic changes that occur as RNAP holoenzyme traverses the various stable intermediates towards elongation complex formation.

## INTRODUCTION

Crp (cAMP receptor protein; also known as CAP, catabolite activator protein) is the most thoroughly characterized transcription factor from a structural and mechanistic standpoint (1-3). It has been the subject of numerous studies focused on unraveling the drivers behind transcription factor activation. These have included, to name a few, comparisons of nuclease protected DNA fragments to elucidate the Crp consensus motif sequence (4-7), mutational analysis of Crp and/or RNA polymerase (RNAP) to reveal the binding interactions that form in distinct promoter architectures (8-15), and three-dimensional structures of Crp and models of it in complex with DNA and RNAP that have been formed (2, 16-19). However, the analysis of Crp and other transcription factors is limited to the *in vitro* model systems for which they are confined and have largely focused on the steps leading to recruitment of RNAP holoenzyme with little attention on the subsequent stages of initiation.

DNA footprinting studies have been instrumental to our understanding of promoter mechanics. This classic approach utilizes the protection from nuclease digestion provided by proteins bound to DNA to produce a highly precise map of the binding site (20). This method has been extensively applied to study the mechanics and kinetics of transcription initiation events (21-23). The outcome of these studies and complementary characterization studies (e.g., x-ray crystallography, single-molecule approaches, and predictive modeling) are at the core of our current, multi-step model of transcription initiation (21-24). However, the rate at which RNAP proceeds through transcription initiation is typically too rapid to be differentiated under physiologically relevant conditions. For example, numerous temperature-modulating experiments have shown that the open RNAP complex dominates at physiological temperatures and that reduced temperatures are needed to recover closed complex intermediates (25-28).

DNA footprinting has also played a significant role in our current understanding of transcription activation by Crp. Detailed *in vitro* studies performed on model promoters (e.g., *lac*, *galP1*, and *deoP2*) have yielded three classes of Crp promoters depending upon the location of the consensus motif sequence(s) relative to the transcription start site (TSS), the number of motif sequences, and the presence of additional transcription factors (1, 2). Class I promoters are thought to mediate activation through a simple recruitment mechanism where interactions are formed between Crp and the α subunit of RNAP yielding the closed promoter complex. Crp forms up to three interactions with RNAP holoenzyme and facilitates isomerization to the open promoter complex at Class II promoters. Class III promoters involve two Crp molecules and a second transcription factor that often represses the activating action of Crp. Footprinting studies under highly controlled and stabilizing conditions have shown that the Crp motif sequence is protected when in complex with Crp and RNAP holoenzyme (29-32). However, these interactions were studied in stabilizing *in vitro* conditions with a focus on characterizing early events during transcription initiation.

Chromatin immunoprecipitation (ChIP) followed by microarray hybridization (chip) or next-generation sequencing have provided genome-scale information on DNA/protein interactions *in vivo*. These techniques have been paramount to studying transcriptional regulators and to construct regulons and transcriptional regulatory networks. However, the information ascertained by application of these methods predominantly provides a binary (present/absent) representation of binding events. Integrating with gene expression analysis allows for expansion of these binary calls to provide conditional activation/repression calls. However, the resolution of ChIP-chip (on the order of kilobases) and ChIP-seq (on the order of hundreds of base pairs) does not enable research to precisely determine the location of the binding event. One of the challenges facing biology is to be able to predict promoter activity. One potential approach to achieve this is by obtaining high-resolution mechanistic information of individual promoters and to convert that mechanistic information into a model of promoter dynamics.

An enhanced form of ChIP-seq called ChIP-exonuclease (ChIP-exo) (33) generates genome-scale maps of DNA binding proteins at single nucleotide resolution. This enables precise identification of binding events by combining DNA footprinting with ChIP. For instance, this method has been applied to the study of eukaryotic pre-initiation complexes, which is typically comprised of RNAP II and no less than six additional general transcription factors (34). The ChIP-exo results were able to spatially resolve individual proteins and agreed strongly with findings produced from crystallographic models. We have previously applied this footprinting assay for application in *Escherichia coli* to elucidate the Fur transcriptional regulon, which predominantly is found to act as a repressor (35).

The study of bacterial transcription activation using high-resolution ChIP-exo data could affirm the transcription initiation processes elucidated *in vitro* under *in vivo* conditions and extend those observations to the genome-scale. Crp provides an ideal entry point for such a study because of the mechanistic and structural information borne out through decades of detailed work on individual promoters (1, 2, 36-39). Here, we applied ChIP-exo to study the DNA protection patterns generated by the housekeeping sigma factor, σ^70^, with respect to published data on RNAP holoenzyme footprinting data. We then compared the protection pattern provided by Crp to σ^70^ and surprisingly found tremendous overlap in their DNA footprinting pattern. However, there was very little observed protection of the Crp motif sequence. This phenomenon was then explored in a repressor, ArcA, and the Crp familial protein, Fnr. Lastly, genetic perturbations to Crp/RNAP interactions were introduced and the affects of these mutations were characterized using ChIP-exo.

## RESULTS

### Strand oriented peak distributions reveal stable intermediates in transcription initiation

The σ^70^ ChIP-exo peak distribution provides the bounds of protected DNA regions on the template and nontemplate strand. ChIP-exo profiles across all binding sites were calculated for both the template and nontemplate strand by first calculating the density of the 5’ end of tags for each individual peak region spanning 400 bp centered and oriented relative to the TSS (transcription start site). The median position of the σ^70^ peak center is 5 bp downstream of the TSS therefore the peak center is found to be an accurate approximation for the TSS (see Supporting Text for detailed discussion). Furthermore, the ChIP-exo profiles for σ^70^ reveal distinctions between the template strand and the non-template strand (Fig. 1A and Fig. S1). The binding profiles show a unimodal distribution on the template strand, whereas a multimodal distribution is seen on the non-template strand. The width of the peak regions was determined by calculating the distance between the maxima on the template and nontemplate strands (Fig. 1B). This indicates that most promoters have a σ^70^ ChIP-exo profile that predominantly fall into one of three groupings.

**Fig. 1.**
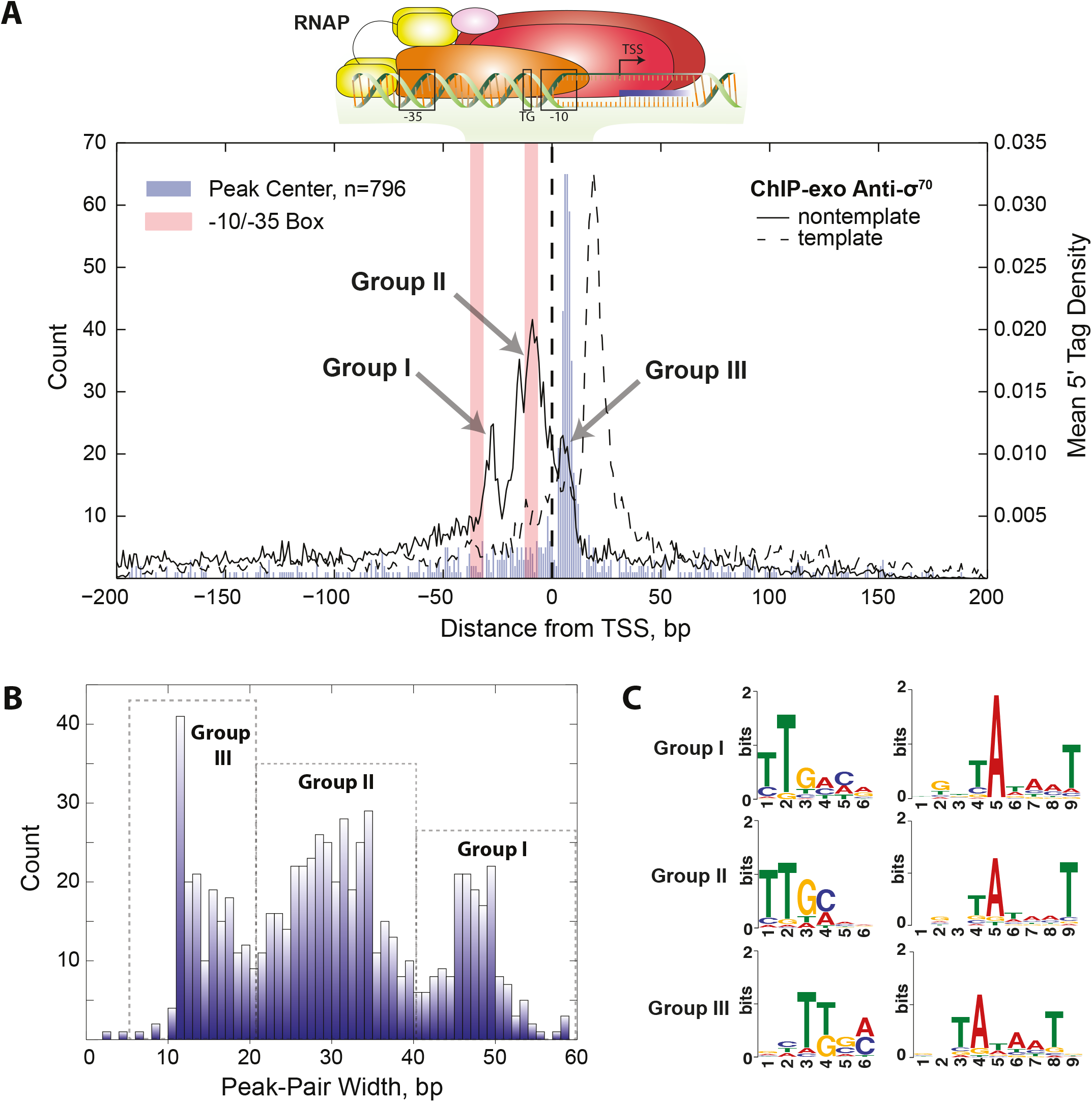
TSS aligned and oriented σ^70^ ChIP-exo data reveals DNA footprint patterns consistent with stable transcription initiation intermediates. (A) ChIP-exo peak regions aligned and oriented relative to the TSS. The peak center (blue bars) is shown to be downstream of the TSS with a median of 5 bp. The mean distribution of the 5’ tags is shown for both strands. The template strand distribution shows a unimodal profile that spans +20±7 bp indicative of RP_O_, ITC, and TEC stable intermediates. The nontemplate strand shows a multimodal distribution with modes centered approximately +5 relative to the TSS (Group III), upstream and over the -10 promoter element (Group II), and slightly downstream of the -35 promoter element (Group I). (B) Examination of the distance between template and nontemplate strand peak maximum shows that the footprint lengths are >40 bp, 21 to 40, <20 and for Group I, Group II, and Group III respectively. (C) A motif search was performed for the -10 and -35 promoter elements for Group I, Group II, and Group III promoters. All three show σ^70^-like promoter sequences with slight differences. Group I has a -35 motif that most closely resembles the consensus (TTGACA), has a highly conserved -11A, and a partial TGn motif. Group III has the least conserved -35 promoter element and no extended -10 promoter element.

The activity of lambda exonuclease is 5’ to 3’ (40) and, as such, the protected region on the template strand is found downstream of the TSS. The unimodal ChIP-exo distribution on the template strand has a maximum 5’ tag density +20 bp downstream of the TSS and approximately 30% of the mean 5’ tag density is found between 20±7 bp. The position of the unimodal distribution on the template strand is in strong agreement with numerous *in vitro* footprinting studies in model promoter constructs characterizing the stable intermediates leading to open complex (RP_O_) formation, the RP_O_, the initial transcribing complex (ITC) and the transition to the ternary elongation complex (TEC). However, the closed promoter complex (RP_C_) does not have an advanced footprint extending to the +20 position (see Supporting Text for detailed discussion).

Unlike the template strand, the ChIP-exo 5’ tag distribution for the nontemplate strand is multimodal. This distribution marks the upstream boundary relative to the TSS. The dominant mode found between -18 and -1 accounts for 28% of the 5’ tag density. Therefore, promoters that belong to this mode have partial to complete protection of the discriminator sequence, the -10 promoter element, and the TGn extended -10 element but little to no protection of the -35 promoter element or any upstream promoter elements (e.g., UP element). The -35 promoter element is partially protected by the mode farthest upstream which accounts for 9% of the 5’ tag density profile and spans -34 to -23 with a maximum located at -28. The upstream boundary, -3, is located in the center of the -35 element. The downstream mode accounts for 8% of the 5’ tag density and is located downstream of the TSS. The boundaries of this mode are between +4 and +12 with a local maximum at +6. Like the template strand, the DNA protected regions of the different modes on the nontemplate strand provide little to no support that recruitment and RP_C_ complex formation is being captured by ChIP (see Supporting Text for detailed discussion).

### Promoter motif analysis of the σ^70^ peak distributions

It is known that promoter sequence elements involved with RNAP holoenzyme recruitment contribute to the post-recruitment kinetics of transcription initiation (22-24). Thus we examined the -10 and -35 promoter elements for the different σ^70^ groups (Fig. 1C) as determined by the difference in peak-pairs (Fig. 1B). σ^70^-like promoter motifs were found in all three groups. Group I, having the longest distance between peak-pairs, has a motif that most resembles the -35 consensus sequence (TTGACA). Furthermore, the -10 promoter element has near perfect consensus at the critical -11A position and a partial TGn motif characteristic of the extended -10 promoter element. Group II resembles the motifs found in Group I but with lower sequence conservation in both the -10 and -35 promoter elements. Conversely, Group III has the most divergent -35 motif from consensus and no appreciable motif for the extended -10 promoter element.

### Promoter characterization of the canonical transcriptional activator, Crp

Transcription factor binding was further studied with ChIP-exo of Crp in *E. coli*. ChIP-exo data showed strong consistency with previously determined Crp binding sites (see Supporting Text). ChIP-exo profiles enabled high-resolution distinction of DNA protection patterns among the three classes of Crp promoters, which are briefly reviewed in the Supporting Text. Representative examples of ChIP-exo profiles generated for cultures exponentially growing in glycerol minimal media (a Crp activating condition) are shown for each of the three Crp Classes (Fig. 2A). The *deoC* promoter is a Class III promoter with two Crp binding sites flanking a CytR regulatory site that represses the activating action of Crp (41). The ChIP-exo protected regions are in close proximity with the three consensus motif sequences with protected regions near -40 and -90 as previously seen *in vitro* (41). However, markedly different profiles are observed in the Class I (*tnaC*) and Class II (*gatY*) promoters that often have no exonuclease protection to the Crp binding site, but instead, have strong protection of the region surrounding the TSS. In fact, these regions correspond greatly with the ChIP-exo profiles generated for σ^70^ under the same condition. However, no observed σ^70^ ChIP-exo peak was detected for the repressed *deoC* promoter.

**Fig. 2.**
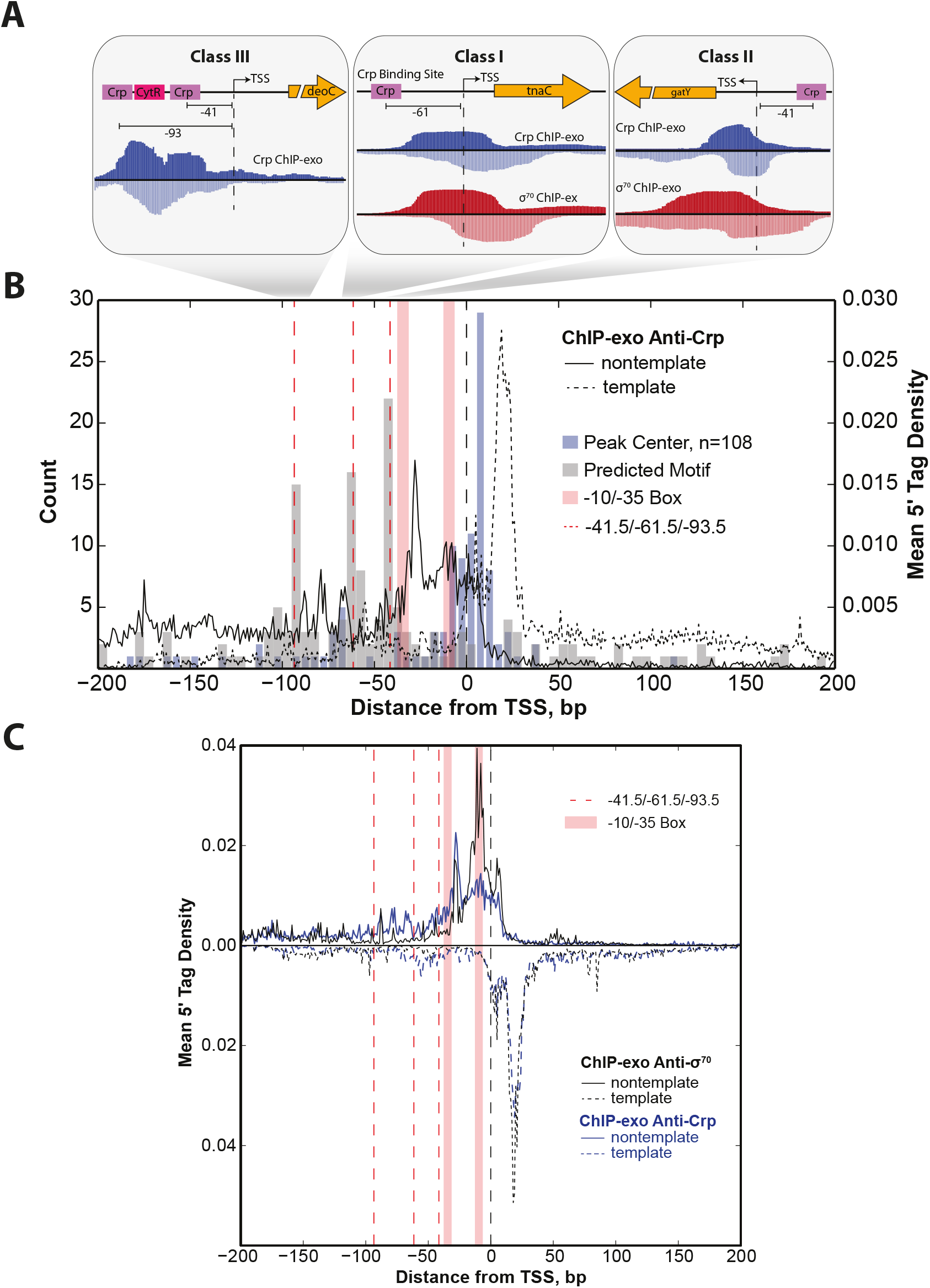
Crp promoter classes have unanticipated ChIP-exo footprint regions. (A) Gene tracks are shown that exemplify the different Crp ChIP-exo footprint profiles observed for the three different classes of Crp promoters. At the Class III promoter *deoC* footprints are found over the Crp motif and the CytR motif which sequesters Crp preventing activation. However, under the activating Class I and Class II promoters there are few observed reads over the Crp motif. Instead, the peak is centered on the TSS and the footprint region co-occurs with that found for σ^70^. Examples of this are shown for *tnaC* (Class I) and *adhE* (Class II). (B) Shown is the mean 5’ tag density ChIP-exo profile aligned and oriented relative to the TSS generated for Crp grown on glycerol minimal media. The distribution of the center position across all Crp peak regions (blue bars) shows close proximity to the TSS. The template strand distribution (dashed black trace) corresponds with the downstream region centered at +20 that is associated with stable intermediates of the RP_O_, the ITC, and the TEC as was observed for σ^70^. The nontemplate strand distribution indicates protection of DNA predominantly occurs downstream of the -35 element with little protection at the predicted binding sites (gray bars). (C) An overlay of the mean 5’ tag density profile of all Crp peak regions (blue traces) and the associated σ^70^ mean 5’ tag density profile in those same peak regions (black traces) illustrates the strong co-occurrence of Crp footprint regions with σ^70^.

The results for these individual promoters are consistent when extended to the genome-scale. Analogous to the analysis performed on σ^70^, all Crp ChIP-exo binding profiles were aligned and strand-oriented relative to the TSS. The same was done with the peak center position and the predicted Crp motif sequence (Fig. 2B). Examination of the motif sites shows three regions of elevated Crp motif sequences centered at -41.5, -61.5, and -93.5 bp upstream of the TSS corresponding with the expected positions of Class II, Class I and Class III promoters respectively (1, 2). However, the mean 5’ tag distribution of Crp ChIP-exo data oriented relative to the TSS illustrates that the peak centers align greatly with the TSS and not the Crp binding site. A similar ChIP-exo profile was obtained when wild type *E. coli* was grown on fructose, another Crp activating condition, but when grown on glucose, a Crp repressing condition, few binding sites were detected and poor alignment was observed relative to the TSS (Fig. S2). We further verified that these results were not artifacts attributed to the anti-Crp antibody used to perform ChIP-exo by generating data on a Δ*crp* strain and no correlation was observed between biological replicate datasets indicating minimal impact due to non-specific binding (Fig. S3). Therefore, the Crp binding profile under activating conditions has poor alignment with the consensus motif sequence.

The ChIP-exo 5’ tag density profile for Crp was also compared with σ^70^ across all Crp binding regions (Fig. 2C). Strand orientated Crp density profiles reveal a unimodal distribution on the template strand and a multimodal distribution on the nontemplate strand analogous to those found for σ^70^. The template strand strongly overlaps the one observed for σ^70^ with a downstream boundary of protected DNA centered on +20 accounting for 33% of the aggregate density profile. However, the Crp nontemplate density profile has distinctive features. First, there is increased DNA protection on the nontemplate strand between the -93.5 and -61.5 markers. This region encompasses 13% of the total 5’ tag density profile. These positions signify the center position of many Class III and Class I Crp motif sequences respectively (1, 2). However, none of these regions indicates protection of the Crp motif sequences found for Class I and Class III promoters and only partial protection for Class II promoters due to the overlap with the -35 box. The strong overlap with the σ^70^ binding profile and alignment with the TSS suggests that Crp immunoprecipitation is occurring in complex with RNAP holoenzyme and, as such, the ChIP profile is more reflective of the stable RNAP intermediates discussed above.

### Rifampicin treated Crp ChIP-exo

Rifampicin (rif) prevents transcription elongation beyond a length of 2-3 nt (42) and, in doing so, leaves the transcription machinery unable to advance beyond the ITC. Therefore, ChIP-exo was performed on cultures treated with rif prior to harvest followed by immunoprecipitation of Crp. The resulting mean 5’ tag density profile generated on both the template and nontemplate strand closely resembles that obtained in the non-rif treated sample (Fig. S4). Therefore, this chemical perturbation of the transcriptional state had no impact on the Crp ChIP-exo distribution and no additional upstream protection of the Crp binding site was observed. This result indicates that the exonuclease footprints are occurring on initiation complexes occurring prior to the TEC. This observation coupled with the evidence against the short-lived RPC complex strongly suggests that the Crp promoters studied here are being captured after dissociation from the motif while they are still bound to RNAP. The capture seems to occur at stable intermediates formed between RPO and the ITC but prior to promoter escape.

### Distinct ChIP-exo profiles for transcriptional activators and repressors

The ChIP-exo binding profiles of activating transcription factors are very different than ChIP-exo profiles of repressing transcription factors. Previous studies have shown transcription factor binding profiles centered on the regulatory motifs in eukaryotic systems (33, 34, 43). Furthermore, we have seen motif centering when ChIP-exo was applied to characterizing the transcriptional repressor Fur in *E. coli* (35). Therefore, we sought to examine if the alignment to the TSS seen in Crp could be extended to the familial protein Fnr and contrasted with the profile generated for a predominantly repressing transcription factor ArcA. ChIP-exo was performed on c-Myc tagged strains of ArcA (repressor) and Fnr (Crp family activator) grown anaerobically on glucose minimal media. The data generated was then processed, aligned, and oriented relative to the nearest TSS (Fig. 3). ArcA, which typically occludes the TSS (44), has no defined ChIP-exo 5’ tag distribution on either strand though there is a noticeable increase in the 5’ tag density around the TSS (Fig. 3A). In contrast, Fnr demonstrates a similar 5’ tag density profile as was seen for Crp and σ^70^ with a strong unimodal distribution on the template strand at +20 and a less defined modal distribution on the nontemplate strand (Fig. 3B). The ArcA ChIP peak regions were aligned relative to the peak center position (Fig. 3C). This resulted in a uniform distribution of 5’ tag density with sharp peaks on the forward (+) strand and the reverse strand (-). Furthermore, plotting the predicted binding sites shows that the protected regions are centered on the ArcA motif. Lastly, the peak-pair differences for ChIP-exo profiles of ArcA and Fnr are shown (Fig. 3D). This reveals that the footprint obtained for the repressor is approximately 30 bp and centered on the binding motif while Crp family activators have a broader footprint distribution centered on the TSS and with a template strand footprint advanced to the +20 position. The broader footprint and advancement to the +20 position affirms the presence of RNAP holoenzyme in the immunoprecipitated complex with little to no protection of the activating motif sequence.

**Fig. 3.**
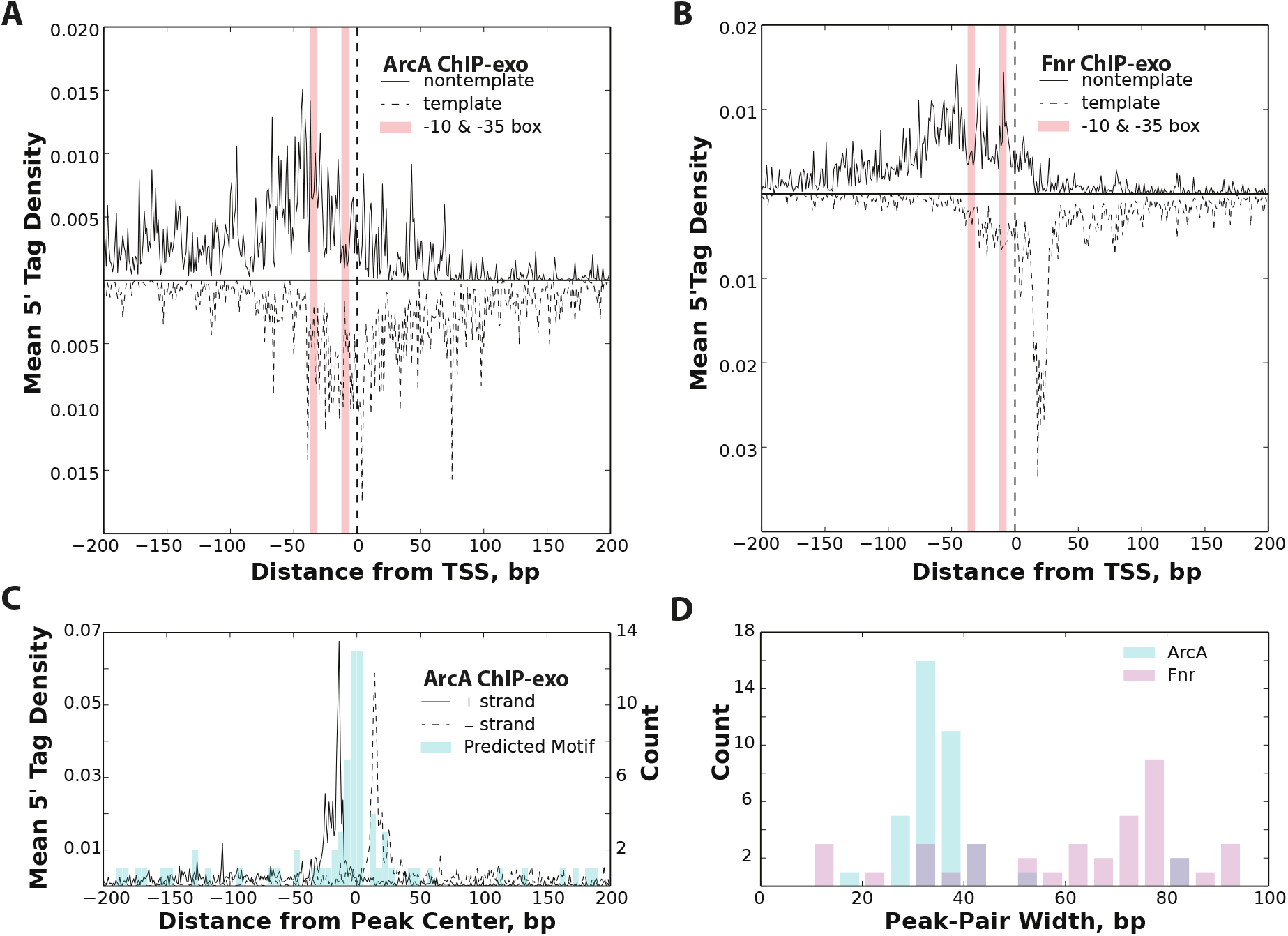
Contrasting ChIP-exo profiles of repressors and activators. (A) The TSS aligned ChIP-exo profile for ArcA, a predominantly repressive transcription factor, is shown to lack the characteristic distribution of mean 5’ tag density observed on both the template and nontemplate strand. (B) The TSS aligned mean 5’ tag density profile for Fnr, typically an activator, resembles the profile found for Crp and ^σ70.^ (C) The ArcA ChIP-exo profile is shown for all peak regions aligned to the peak center position. Also shown is a histogram of the center of the predicted ArcA binding site relative to the peak center position. This illustrates that the ChIP-exo profile is centered on the predicted binding site. (D) A comparison of the peak-pair distance is shown to illustrate the difference in resolution observed between ArcA and Fnr. ArcA, the repressor, is revealed to have shorter footprints compared with Fnr, the activator.

### Genetic perturbation of RNAP holoenzyme/Crp interactions

We next sought to determine the impact of genetic perturbations to the RNAP holoenzyme/Crp interactions by introducing deleterious mutations to the Activating Regions (see Supporting Text for discussion) Ar1 and Ar2. Mutations were introduced to create ΔAr1, ΔAr2, and ΔAr1ΔAr2 mutants (Fig. 4A). ChIP-exo was performed on these mutant strains with glycerol as the sole carbon source. In comparison with the wild type, each mutant resulted in the loss of peak regions (Fig. 4B). The most drastic effect was observed in the ΔAr1ΔAr2 mutant which retained ~40% of the peaks in the wild type strain. This result indicates the importance of these Ar interactions on the stabilization of both Crp and RNAP holoenzyme at the promoter site. Furthermore, the characteristic ChIP-exo 5’ tag density profiles (see Fig. 2C) on both strands were systematically degraded with each mutation resulting in profiles that no longer aligned well to the TSS (Fig. S5). To determine which peak regions were lost as a result of these genetic perturbations, the distribution of peak region centers was analyzed (Fig. 4C). The mutations predominantly result in a loss of peak-regions where the peak center was located near the TSS (-10 to +20 bp) and peak centers farther away from the TSS were less impacted. Lastly, the distribution of predicted binding sites were examined in the context of the different mutant strains (Fig. 4D). In agreement with expectation, modulation of Ar1 results in a drop in the predicted binding sites observed near -61.5, the typical Class I promoter distance from the TSS. This drop near -61.5 was partially recovered in the Ar2 mutant but a severe drop in the -41.5 centered binding sites occurred. This distance upstream of the TSS is associated with Class II promoters. The ΔAr1ΔAr2 mutant has a loss in peak regions with Crp binding sites matching those of Class I and Class II promoters. However, the peak regions of Class III found near the -93.5 position are unaffected by mutations in Ar1, Ar2, or both.

**Fig. 4.**
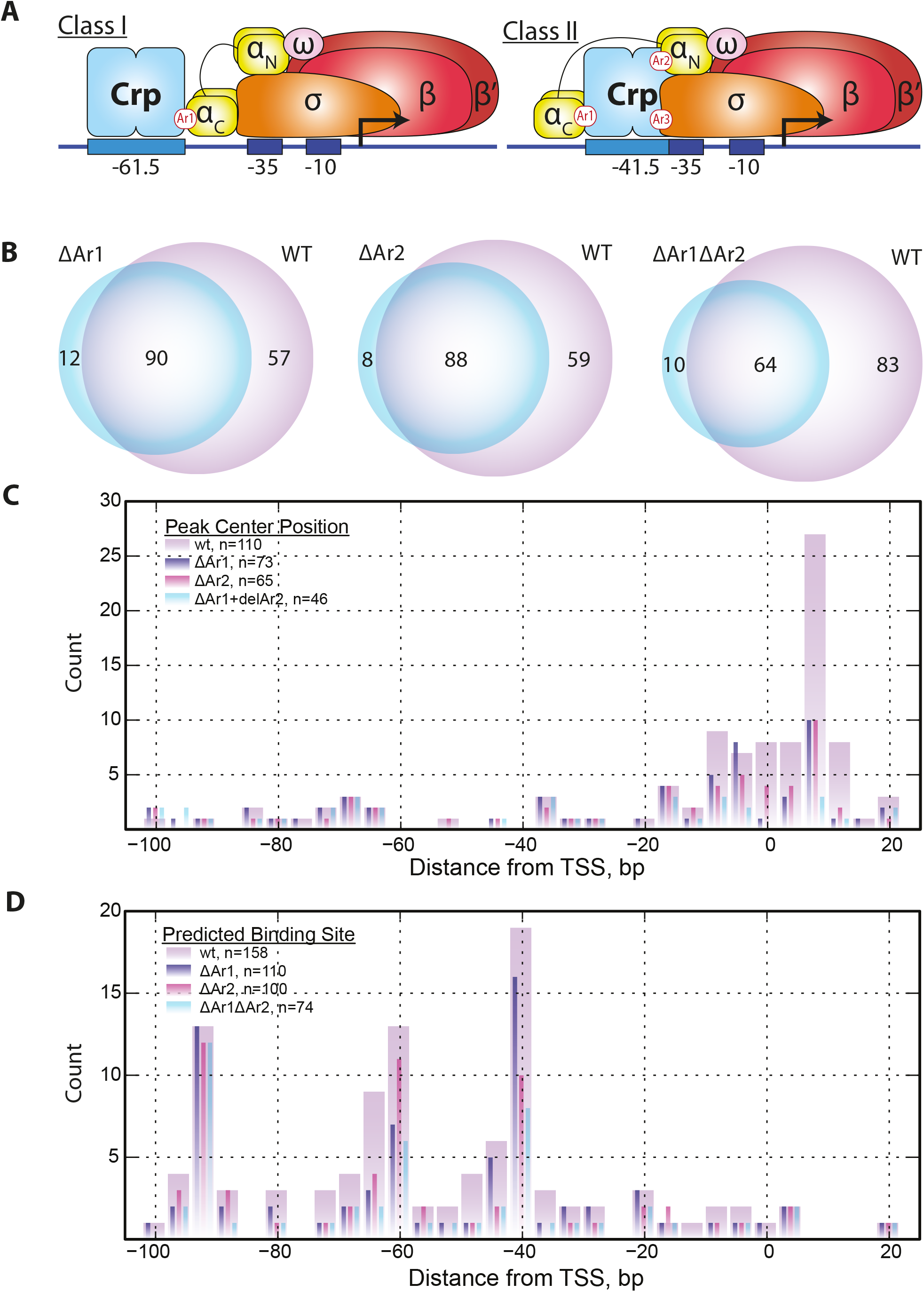
The effect of genetic perturbation on Crp/RNAP interactions. (A) Cartoon illustrating the interactions between activating regions (Ar’s) and RNAP for Class I and Class II activators. Crp Class I promoters make a single contact with RNAP at Ar1 whereas Crp Class II activators make upwards of three contacts (Ar1, Ar2, and Ar3). Deletions of Ar1, Ar2, and Ar1+Ar2 where generated. (B) Venn diagram showing pairwise comparison of peaks regions detected for ΔAr1, ΔAr2, and ΔAr1ΔAr2 with wild type Crp. All cultures were grown with glycerol as the carbon source. The mutations to Crp result in fewer detected peaks relative to wild type Crp indicating promoter destabilization. (C) Histogram of the peak center position relative to the TSS for wild type Crp, ΔAr1, ΔAr2, and ΔAr1ΔAr2 mutants. This illustrates that the peak centers nearest the TSS (-15 to +20) are predominantly affected by deletion of Ar1 and Ar2, whereas peak regions centered upstream of the TSS (< -15) are largely unaffected. (D) An alternative view to the histogram shown in (C) that shows the distribution of predicted Crp binding sites relative to the TSS. The ΔAr1 strain shows a reduction in the number of peak regions with -61.5 motifs compared with wild type and ΔAr2 indicating a sensitivity of Class I promoters to mutations to this region. Similarly, the ΔAr2 strain shows a substantial loss of Class II associated peak regions (-41.5 binding sites) compared with Class I (-61.5). The ΔAr1ΔAr2 mutant shows reductions in both -41.5 and -61.5 binding sites compared with the wild type. None of the ΔAr strains showed a reduction in the peak regions with Class III binding sites (e.g., -93.5 binding sites).

## DISCUSSION

Here we present high-resolution ChIP-exo datasets that enable *in vivo* characterization of transcription initiation events at the genome-scale. The detailed footprinting performed on *E. coli* σ^70^ are foundational to subsequent analysis of transcription activation associated with Crp family proteins. The σ^70^ ChIP-exo profiles reflect findings determined in a number of *in vitro* footprinting studies performed on individual model promoters (for detailed discussion see Supporting Text). Those studies have revealed that shortly after recruitment, the RNAP holoenzyme complex advances to the +20 position relative to the TSS. The upstream footprint boundary is less pronounced following an oscillatory pattern covering different promoter elements. In strong agreement with what *in vitro* studies have revealed, the σ^70^ ChIP-exo data presented in Fig. 1 shows the template strand DNA boundary located at the +20 advanced position. Similarly, the nontemplate strand data shows a multimodal distribution that poorly protects the -35 and promoter elements upstream thereof. Thus, comparison of the σ^70^ ChIP-exo data to *in vitro* footprinting profiles of RNAP holoenzyme indicate that the ChIP results generated here, and likely elsewhere, are recovering the entire RNAP holoenzyme complex. While this may not be a surprising result, the comparison also determines that RNAP holoenzyme complex is most often captured after recruitment to the promoter. The +20 advancement on the template strand is characteristic of stable RNAP holozenyme initiation intermediates that occur post-recruitment. The advanced +20 position has been observed for stable RPC intermediates, RPO, ITC, and early TEC complexes but not for the recruited RNAP holoenzyme complex whose footprint does not extend far beyond the TSS. Given the time scale of ChIP crosslinking is on the order of minutes, it is likely that ChIP studies characterize RNAP holoenzyme at kinetically long-lived, stable states formed on the path towards a promoter-escaped, elongation complex. *In vitro* kinetic studies support the ChIP-exo data presented here where the rate-limiting step during transcription initiation is most often downstream of recruitment. Genome-scale characterization studies of bacterial promoters have also determined that the rate-limiting step in transcription predominantly occurs post-recruitment of RNAP (45-47). Therefor, the σ^70^ ChIP-exo results affirm *in vitro* results observed in model promoter systems and extend those findings to the genome-scale and under *in vivo* conditions.

Surprisingly, the observations made for σ^70^ were also observed in datasets where anti-Crp antibodies were used to study the binding patterns of this well characterized transcription factor. Crp binding profiles did not align to the motif sequence as would be expected but, rather, were centered on the TSS. These binding profiles, like σ^70^, largely exhibit advancement of the DNA protected boundary to the +20 position on the template strand. Furthermore, the binding pattern on the nontemplate strand shows little to no protection of the well-characterized Crp binding motifs. These results indicate that Crp and RNAP holoenzyme are not only co-immunoprecipitated during ChIP experiments, but also that the subsequent Crp ChIP-exo footprint patterns reflect the same long-lived RNAP holoenzyme transcription initiation intermediates observed with σ^70^. Though this study cannot definitively rule out that this observation can be attributed to limitations of formaldehyde crosslinking, several pieces of supporting information suggest otherwise. First, the same binding pattern observed in ChIP-exo studies performed on the native *crp* gene using an anti-Crp antibody were also observed in a c-*myc*-tagged *crp* gene fusion strain of *E. coli* K12 using an anti-c-myc antibody. Second, the closely related transcription factor, Fnr, yielded analogous ChIP-exo profiles to Crp on the template and nontemplate strands whereas the ChIP-exo profile of ArcA, a predominantly repressing transcription factor (48), showed a completely different binding profile. ArcA, as well as previously published work on Fur (35), demonstrate strong centering on the DNA motif and a narrower footprint compared with Crp and Fnr. Third, systematic mutations disrupting Crp/RNAP interactions revealed a significant loss in ChIP signal in Class I and Class II activating promoters but little disruption to the Class III promoters. Therefore, ChIP-exo binding sites associated with RNAP were eliminated in Crp-RNAP binding deficient mutants, whereas, binding events not associated with RNAP, namely those distant from the TSS, were still observed and motif centered. In fact, all conditions tested showed a subset of Crp binding sites that are motif centered. Thus, the alignment of ChIP-exo data relative to the TSS and the advancement of the template strand ChIP-exo distribution to the +20 position appear to be characteristic of transcriptional activation, whereas peak regions aligning relative to the motif sequence are not. Nevertheless, subsequent orthogonal confirmation of observations resulting from ChIP-exo studies would be beneficial though doing so under *in vivo* conditions and at the genome-scale is currently not feasible.

The vast majority of *in vitro* Crp studies have focused on the mechanism of recruitment and this transcription factor’s role in transcription initiation. However, a series of experiments were performed that deciphered the role of Crp in transcription after the RP_O_ complex was formed. A heparin challenge was applied to different Crp promoter classes to displace Crp from the open ternary complex (49, 50). In every promoter characterized, removal of Crp was inconsequential to transcriptional output upon open complex formation. These studies established that Crp plays a role in the recruitment of RNAP holoenzyme and also the isomerization of RNAP holoenzyme to form the open complex (1, 2). Thereafter, Crp’s presence or absence at the DNA binding site has no impact on transcription. Therefore, it is plausible that as RNAP holoenzyme transverses through the post-recruitment stages of transcription initiation, Crp is displaced from the DNA binding motif but remains bound to RNAP holoenzyme until promoter escape (Fig. S6). This hypothesis would explain the Crp ChIP-exo footprinting pattern that closely resembles that of RNAP holoenzyme and the poor protection of Class I and Class II DNA motif sequences. In addition to the data discussed above, Crp binding profiles in the presence of rifampicin indicate that Crp remains bound to the RNAP holoenzyme up to and including TEC formation (Fig. S4).

However, the data generated in this study alone cannot resolve what drives Crp/DNA dissociation to occur or how the release of Crp occurs from RNAP holoenzyme. The mechanisms driving σ factor release have proven to be elusive (21, 51-53) and the release of transcriptional activators will likely be just as elusive. It is thought that the energy needed for promoter escape is established through a stressed intermediate resulting from scrunching (54, 55). This stressed intermediate may break the bonds between the σ factor and RNAP enabling RNAP to proceed to the elongation stage of transcription while the σ factor is retained at the promoter or dislodged from the promoter. Perhaps scrunching provides sufficient energy to also break the bonds formed between Crp, the σ factor, and RNAP, thereby enabling full transition into transcription elongation.

The detailed molecular interactions elucidated here reflect transitions of RNAP during transcription initiation at the genome-scale. This study is merely a starting point with numerous potential applications for ChIP-exo in studying promoter dynamics. The challenge will be to integrate multi-scale approaches such that we advance beyond studying just binary interactions of transcriptional regulators and begin to quantitatively unravel the molecular dynamics of transcription initiation. We believe that the datasets and analytical approaches utilized here provide a key component towards possibly reconstructing a quantitative, mechanistic, predictive model of promoter dynamics at the genome-scale.

## Materials and Methods

### Strains and Culturing Conditions

*Escherichia coli* MG1655 cells and derivatives thereof were used for all experiments. Fnr-8-myc, and ArcA-8-myc tagged strains were previously constructed (56). The Δ*crp* strain was generated by replacing native gene with a kanamycin resistance marker from start codon to stop codon using the λ red mediated gene replacement method described (57). The Δ*crp* was used as a basis for constructing the ΔAr1, ΔAr2 and ΔAr1ΔAr2 mutant strains using a modification of the λ red mediated gene replacement method. Briefly, plasmids carrying the different Ar mutant sequences were *de novo* synthesized using GeneArt (Life Technologies) with restriction sites at the 5’ and 3’ end of the gene. The gene was digested from GeneArt plasmids and ligated into the pKD3 plasmid directly upstream of the chloramphenicol (Cm) resistance gene. Resulting plasmids have the Ar mutant-*crp* gene, followed by the FRT flanked Cm resistance cassette as in pKD3 plasmid. Linear PCR products were amplified from resulting modified pKD3 plasmids using primers with 5’ overhangs with homology directly upstream of the start codon and downstream of the stop codon of *crp* gene to direct the insertion. This PCR product was transformed into electrocompetent Δ *crp E. coli* K12 carrying the pKD46 plasmid, and selected by Cm resistance, correct insertions were verified by Sanger sequencing. The Cm resistance gene was then removed from confirmed mutant strains by FLP recombinase excision transforming with pCP20 plasmid as previously described (57). The ΔAr1 mutant introduces a mutation to the Ar1 region, HL159, previously determined to break contacts between Ar1 and the α subunit of RNAP (13, 58). The ΔAr2 mutant does the same for Ar2 but introduces two mutations, KE101 and HY19 (58). The ΔAr1ΔAr2 strain carries the HL159 mutation and the KE101 mutation.

M9 minimal media was used for all cultures with 2 g/L of glucose, fructose, or glycerol. For σ^70^, Crp, Δ*crp*, ΔAr1, ΔAr2, and ΔAr1ΔAr2 experiments, cultures were grown aerobically in shake flasks. Rifampicin conditions were incubated in the presence of rifampicin (50 μg/mL final concentration) for 20 min prior to crosslinking as previously described (59). Fnr and ArcA experiments were conducted similarly but grown under anaerobic conditions.

### ChIP-exo Experiments

The ChIP-exo protocol was adapted from Rhee and et al. for the Illumina platforms with the following modifications (33). DNA crosslinking, fragmentation, and immunoprecipitation were performed as previously described (60) unless otherwise stated. Clarified lysate was continuously sonicated at 4 °C using a sonicator bell (6W) for 30 min. Antibodies used in this study are: anti-Crp (Neoclone N0004), anti-σ^70^ (Neoclone WP004), and anti-Myc (Santa Cruz Biotechnology sc-40). Immunocomplexes were captured using Pan Mouse IgG Dynabeads (Life Technologies). The following library preparation steps were sequentially performed while the protein/DNA/antibody complexes were bound to the magnetic beads: end repair (NEB End Repair Module), dA tailing (NEB dA-Tailing Module), adaptor 2 ligation (NEB Quick Ligase), nick repair (NEB PreCR Repair Mix), lambda exonuclease treatment (NEB), and RecJf exonuclease treatment (NEB). A series of step-down washes were done between all steps using buffers previously described (60). Strand regeneration and library preparation followed the approach of Rhee et al. with the exception of a 3’ overhang removal step after the first adaptor ligation and prior to PCR enrichment by treating with T4 DNA Polymerase for 20 min at 12 °C. Libraries were sequenced on an Illumina MiSeq. Reads were aligned to the NC_000913.2 genome using bowtie2 (61) with default settings. Peak calling was performed using GPS in the GEMS analysis package (62) with the ChIP-exo default read distribution file with the following parameter settings: mrc 20, smooth 3, no read filtering, and no filter predicted events. GPS was used over GEMS because GEMS peak boundaries are influenced by motif identification whereas GPS is not. ChIP-peak calls were manually curated for anti-Crp (wt and Ar mutant strains) and anti-Myc (Fnr, and ArcA) for all substrates and conditions. A superset of GPS peak calls across all anti-Crp conditions was analyzed for presence/absence in each individual condition.

### Gene Expression

Gene expression analysis was performed using a strand-specific, paired-end RNA-seq protocol using the dUTP method (63). Total RNA was isolated and purified using the Qiagen Rneasy Kit with on-column DNase treatment. Total RNA was depleted of ribosomal RNAs using Epicentre’s RiboZero rRNA removal kit. rRNA depleted RNA was then primed using random hexamers and reverse transcribed using SuperScript III (Life Technologies). Sequencing was performed on an Illumina MiSeq. Reads were mapped to the NC_000913.2 reference genome using the default settings in bowtie2 (61). Datasets were quantified using cuffdiff in the cufflinks package to generate FPKM (Framents Per Kilobase per Million reads mapped) values for all genes (64).

### Data Deposition

Datasets are located at the Gene Expression Omnibus under Accession number GSE64849. Reviewer link: http://www.ncbi.nlm.nih.gov/geo/query/acc.cgi?token=cdunciiszdkzpob&acc=GSE64849

## AUTHOR CONTRIBUTIONS

H.L. and S.F. conceived and planned the work presented here. H.L., J.T., and R.S. performed all of the experiments. H.L. and R.S. adapted the ChIP-exo protocol for microbial applications. S.F., H.L., and A.E. processed and analyzed all of the data. J.U. and K.Z. provided guidance for experimental design. H.L., S.F., A.E., and B.O.P. wrote and edited the manuscript.

## ACKNOWLEDGEMENTS

The Novo Nordisk Foundation and NIH Grants GM057089 and GM102098 provided financial support for this work. H.L. was supported through the National Science Foundation Graduate Research Fellowship under grant DGE1144086.

